# Brain age prediction and deviations from normative trajectories in the neonatal connectome

**DOI:** 10.1101/2024.04.23.590811

**Authors:** Huili Sun, Saloni Mehta, Milana Khaitova, Bin Cheng, Xuejun Hao, Marisa Spann, Dustin Scheinost

## Abstract

Structural and functional connectomes undergo rapid changes during the third trimester and the first month of postnatal life. Despite progress, our understanding of the developmental trajectories of the connectome in the perinatal period remains incomplete. Brain age prediction uses machine learning to estimate the brain’s maturity relative to normative data. The difference between the individual’s predicted and chronological age—or brain age gap (BAG)—represents the deviation from these normative trajectories. Here, we assess brain age prediction and BAGs using structural and functional connectomes for infants in the first month of life. We used resting-state fMRI and DTI data from 611 infants (174 preterm; 437 term) from the Developing Human Connectome Project (dHCP) and connectome-based predictive modeling to predict postmenstrual age (PMA). Structural and functional connectomes accurately predicted PMA for term and preterm infants. Predicted ages from each modality were correlated. At the network level, nearly all canonical brain networks—even putatively later developing ones—generated accurate PMA prediction. Additionally, BAGs were associated with perinatal exposures and toddler behavioral outcomes. Overall, our results underscore the importance of normative modeling and deviations from these models during the perinatal period.

## Introduction

The human connectome—a whole-brain connectivity map—undergoes rapid changes during the perinatal period. The structural connectome is established across the third trimester^1^. For example, hallmark topological properties of the structural connectome, including rich-club connections^2^, network controllability^3^, and connectome fingerprinting^4^, are observable at birth. Structural connections mature further during the first year of life but slow down afterward^5^. In contrast, the functional connectome develops slower in the fetal period. Early functional connections are observed in the fetal period^6^, and lower-order networks (e.g., motor-sensory) are present at birth^5^. However, many functional networks (e.g., the default mode network) only resemble older children after the first year of life^7,8^. Nevertheless, emerging research suggests that most functional networks are detectable in the neonatal period^9^ with larger samples and higher-quality data. Thus, despite progress in mapping developmental trajectories in the perinatal period, a normative model of the early connectome does not exist.

Brain age prediction is a novel approach to normative modeling^10,11^. This approach consists of two steps. First, a machine learning model is trained to predict an individual’s age using neuroimaging data. These predicted ages represent the brain’s maturity relative to the normative data. Predicted ages for different modalities can be compared to investigate correlational patterns of developmental trajectories across the modalities. For example, a strong correlation between brain age from structural and functional connectomes suggests synchronized structural and functional development. Second, the difference between the individual’s predicted and chronological age—or brain age gap (BAG)—is calculated. BAGs represent the deviation from normative brain measures at any particular age. Other participant characteristics (e.g., behavior or disease symptoms) can be associated with BAGs to investigate their impact on normative development or aging. While research using brain age predictions and BAGs is robust in adolescents and adults^12,13^, they are not well-studied with connectome data from the perinatal period. Four studies^9,14–16^ have used functional connectomes, and two^17,18^ have used structural connectomes to predict age in the first months of life. However, none of these studies have compared structural and functional brain ages.

Early life exposures and later toddler behavior are important phenotypes to investigate with normative developmental models, like BAGs. Numerous perinatal exposures, including maternal mental health^19–22^, physical health^23–25^, and substance use^26,27^, contribute to individual differences in neurodevelopmental trajectories. These exposures push and pull development to better or worse trajectories. Likewise, early deviations from a normative model may reflect future risk of delays or disorders. Therefore, BAGs during the perinatal period might eventually help with risk identification. Previous studies have not correlated BAGs with early life exposures. Further, only one correlated BAGS with later toddler behavior in preterm infants^17^.

Here, we assess brain age prediction and BAGs of structural and functional connectomes for infants in the first month of life. We asked the following questions: How do structural and functional connectomes mature during the perinatal period? How correlated is structural and functional development at this age? How do individual canonical brain networks mature in distinctive patterns after birth? How do perinatal maternal exposures alter normative development (e.g., BAGs)? How do BAGs in infants predict toddler cognition and behavior? To answer these questions (Fig. 1), we calculated brain age and BAGs based on resting-state fMRI and DTI for 611 infants (174 preterms; 437 terms) from the Developing Human Connectome Project^28^ (dHCP, http://www.developingconnectome.org/) using connectome-based predictive modeling (CPM). Both structural and functional connectomes accurately predicted postmenstrual age (PMA) for preterm and term infants. Predicted PMA from structural and functional connectomes were correlated, suggesting a coupling between structural and functional development. Several canonical brain networks predicted PMA. Multiple prenatal maternal effects altered BAGs in the two groups, and BAGs were associated with development outcomes in toddlerhood. Together, we established normative models of connectome development in infants and highlighted that deviations associate with early-life maternal exposures and later behavior.

**Fig. 1.**
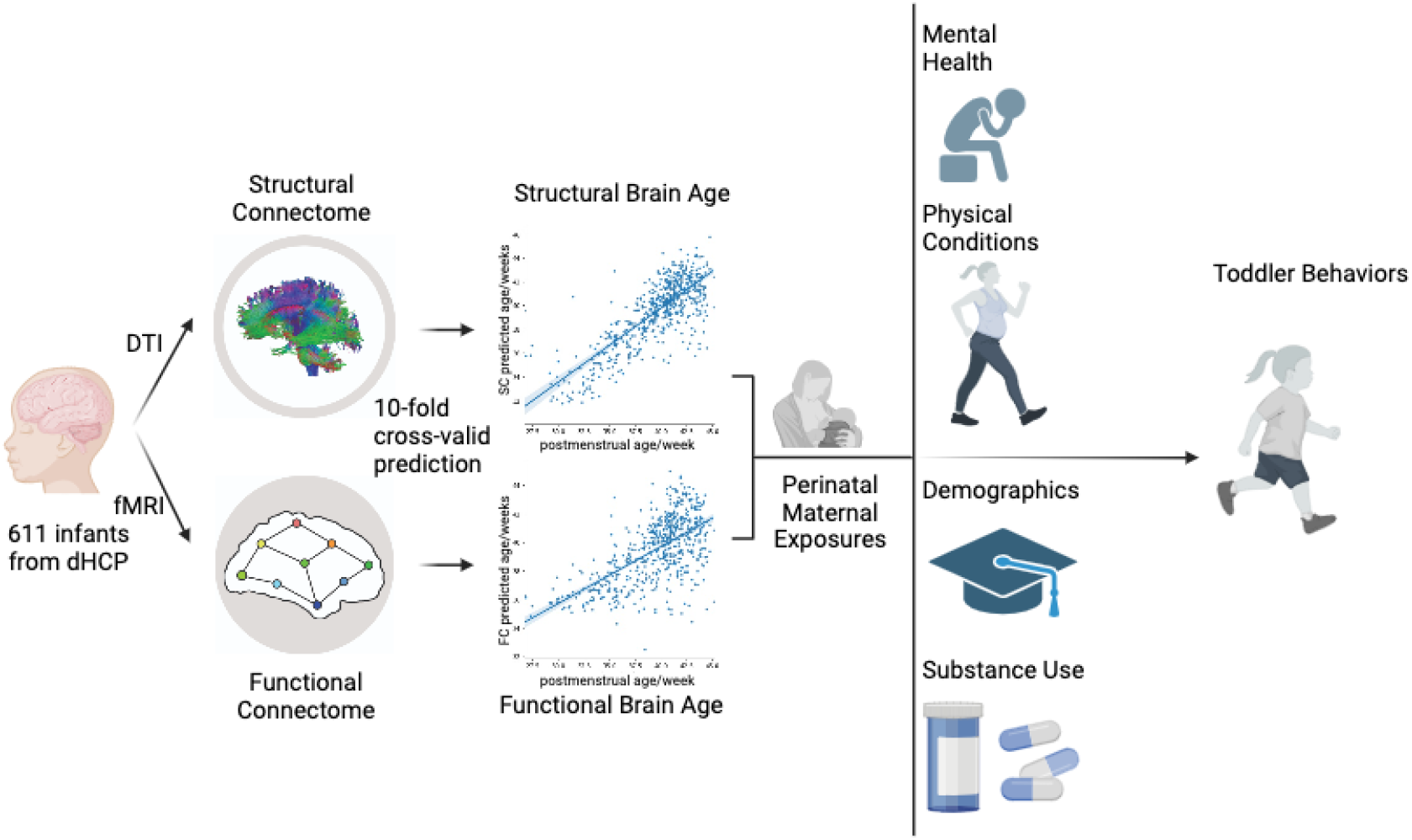
Overview of study design. Predictive models for infants’ postmenstrual ages were built using structural and functional connectomes separately for term and preterm infants. Brain age gaps (BAGs) were calculated as the difference between predicted and actual postmenstrual age. Structural and functional BAGs were associated with perinatal effects and later cognitive behaviors.

**Fig. 2.**
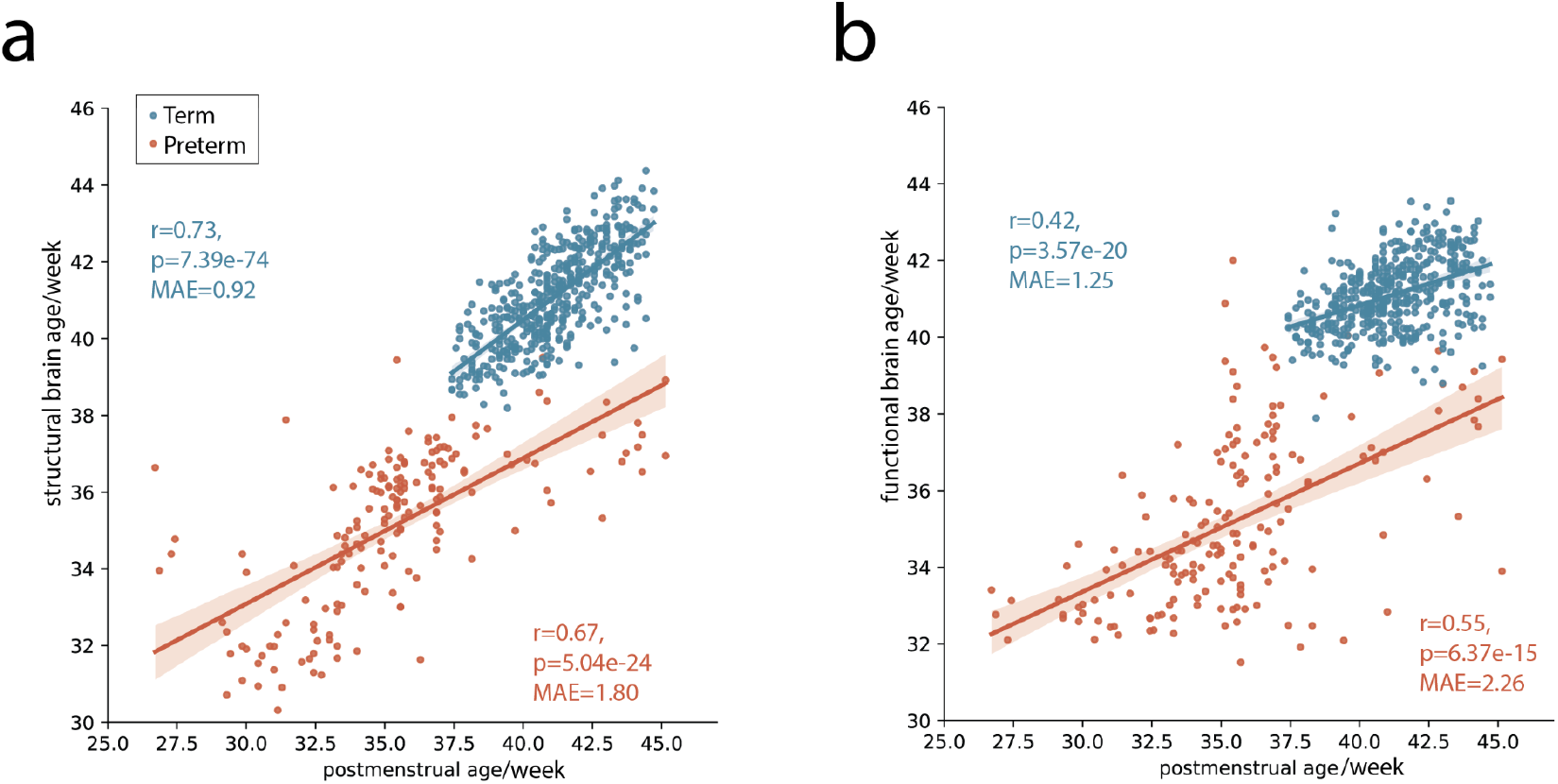
Structural and functional connectomes predict postmenstrual age (PMA) in term and preterm infants. **(a)** PMA was accurately predicted based on structural connectomes for term (blue dots, r=0.73, p=7.39e-74) and preterm infants (red dots, r=0.67, p=5.04e-24). **(b)** PMA was accurately predicted based on function connectomes for term (r=0.42, p=3.57e-20) and preterm infants (r=0.55, p=6.37e-15). Models from structural connectomes were more accurate than models from functional connectomes (z-test, one-tail; term: z=7.96, p=4.32e-14; preterm: z=2.16, p=0.03).

## Results

We used multimodal neuroimaging data for 611 infants (174 preterms; 437 terms) from the developing Human Connectome Project (dHCP). Term infants were born between 37.00 and 42.29 weeks of gestation and scanned between 37.43 and 44.71 weeks. Preterm infants were born between 23.00 and 36.86 weeks of gestation and scanned between 26.71 and 45.14 weeks. Demographic information about the infants is presented in Table 1. Functional connectomes were created from resting-state functional magnetic resonance imaging (fMRI) data. Structural connectomes were created from diffusion weighted imaging (DWI) data. We predict postmenstrual ages (PMA) using connectome-based predictive modeling (CPM) with 10-fold cross-validation. All models controlled for sex, brain volume, and head motion. Separate models were built from the functional and structural connectomes for the term and preterm infants. Results were replicated using an additional machine learning method (support vector regression) and shown in the Supplement (Table S4-6).

**Table 1.** Infant characteristics, mean (std)

|  | preterm(n=174) | term (n=437) | *p* Value |
| --- | --- | --- | --- |
| Sex, num of females (%) | 74 (43.53%) | 203 (46.45%) | .38 |
| Gestational age (GA) / weeks | 31.95 (3.72) | 39.90 (1.27) |  |
| Post Menstrual Age(PMA) / weeks | 35.36 (3.58) | 41.14 (1.72) |  |
| Bayley Scales Of Infant and Toddler Development (BSID) | n=129 | n=345 |  |
| Cognition | 9.74 (2.53) | 10.11 (2.09) | .10 |
| Language | 18.44 (5.80) | 19.26 (5.09) | .13 |
| Motor | 19.67 (3.71) | 20.62 (3.16) | < 0.01 |
| Child Behavior Checklist (CBCL) | n=125 | n=337 |  |
| internalizing | 45.62 (10.55) | 44.41 (9.61) | .24 |
| externalizing | 48.54 (9.50) | 47.53 (9.23) | .30 |
| total | 47.90 (9.86) | 46.91 (9.22) | .32 |
| Early Childhood Behavior Questionnaire (ECBQ) | n=124 | n=338 |  |
| surgency | 5.25 (0/90) | 5.20 (0.73) | .56 |
| negative affective | 2.58 (0.78) | 2.63 (0.67) | .46 |
| effective control | 4.59 (0.69) | 4.53 (0.71) | .54 |
| Quantitative Checklist for Autism in Toddlers (Q-CHAT) | n=125 | n=338 |  |
| total | 30.48 (10.30) | 30.08 (8.82) | .68 |

## Structural and functional connectome predict postmenstrual age in infants

Using structural connectomes, we significantly predicted PMA for term (r=0.73, p=7.39e-74; mean absolute error (MAE)=0.92 weeks) and preterm infants (r=0.67, p=5.04e-24; MAE=1.80). Similarly, using functional connectomes, we significantly predicted PMA for term (r=0.42, p=3.57e-20; MAE=1.25) and preterm infants (r=0.55, p=6.37e-15; MAE=2.26). Models from structural connectomes were more accurate than models from functional connectomes (z-test, one-tail; term: z=7.96, p=4.32e-14; preterm: z=2.16, p=0.03), suggesting a larger association between structural development and PMA during early infancy. Predicted ages from the structural connectomes (i.e., structural age) significantly correlated with predicted ages from the functional connectomes (i.e., functional age; term: r=0.36, p=4.11e-15; preterm: r=0.50, p=2.32e-12), suggesting a moderate structural and functional age coupling. This correlation was stronger in the preterm infants than in term infants (z=-1.91, p= 0.028). Finally, BAGs from the structural connectomes significantly correlated with BAGs from the functional connectomes for preterm (r=0.22, p=0.0044) but not term infants (r=0.092, p=0.054) when controlling for PMA.

## Structural and functional brain age for canonical networks

We investigated how connections within and between canonical brain networks predicted PMA to provide PMA prediction at greater spatial specificity. Connectomes were divided into eight canonical brain networks: visual, somatomotor, frontoparietal, default mode, dorsal attention, ventral attention, limbic, and subcortical networks. For each network, the predicted age was generated based only on the connections within this network (i.e., connections between nodes in the same network). For structural and functional connectomes, all but the ventral attention network predicted PMA for term infants. In both cases, the frontoparietal network showed the highest correlation between predicted and observed PMA (structure: r=0.59, p=6.05e-43, MAE=1.12 weeks; function: r=0.28, p=1.41e-09, MAE=1.41 weeks). Like the whole-brain models, structure better predicted PMA than function in all networks (z’s>2.58, p’s<0.0097) except for the dorsal attention network (z=-0.47, p=0.63). In contrast, function predicted PMA better than structure for preterm infants. All but the dorsal and ventral attention networks from the functional connectome significantly predicted PMA in the preterm infant group. Only the limbic, default mode, and subcortical networks from the structural connectome predicted the PMA preterm infants. Models generated from a single network were numerically worse than the whole-brain models.

Structural and functional age for each network were correlated except for the dorsal attention network in term infants. Nevertheless, these correlations (r’s<0.18) were smaller than at the whole-brain level (r=0.36). For the three networks exhibiting significant age predictions for both structural and functional connectomes, significant correlations between structural and functional ages were observed for preterm infants (r’s>0.25, p’s<0.001). Structural and functional ages were more correlated in the preterm infants than in the term infants for the default mode (z=-2.82, p=0.0024), the subcortical (z=-3.1907, p=0.00071), but not the limbic networks (z=-1.27, p=0.10).

Predictions using between-network connections are shown in Fig. S1. For structural connectomes, all between-network models, except for connections between the dorsal and ventral attention networks, predicted PMA for the term infants. For structural connectomes, between-network models predicted better (on average) than within-network models (between-network: mean r=0.54; within-network: mean r=0.45; p=0.046, permutation testing, 1000 iterations). Like other analyses in term infants, between-networks models from structural connectomes performed better than models from functional connectomes. No differences in performance for within- and between-network models were observed for the functional connectomes. Similar trends were observed for models in preterm infants. Finally, we repeated PMA prediction based on individual lobes from structural connectomes. These results are shown in Fig. S2.

After calculating each network’s structural and functional ages, we correlate the structural and functional ages between different canonical networks. A large correlation between the two networks’ structural (or functional) age suggests that these networks have synchronous development. For term infants, structural ages were significantly correlated (r’s>0.13, p’s<0.05) across networks except for the structural ages from the dorsal attention and subcortical networks (Fig. 3C). As for functional ages, most were correlated across networks, except the functional ages from the frontoparietal and subcortical networks and the functional ages from the visual network to the other networks (Fig. 3C). Together, these results suggest that networks develop in synchrony in the term infant group. For preterm infants, all networks that predict PMA show strong correlations (r’s>0.53, p’s<0.001) in structural and functional ages between the different networks. These correlations were generally stronger in preterm infants than in term infants.

**Fig. 3.**
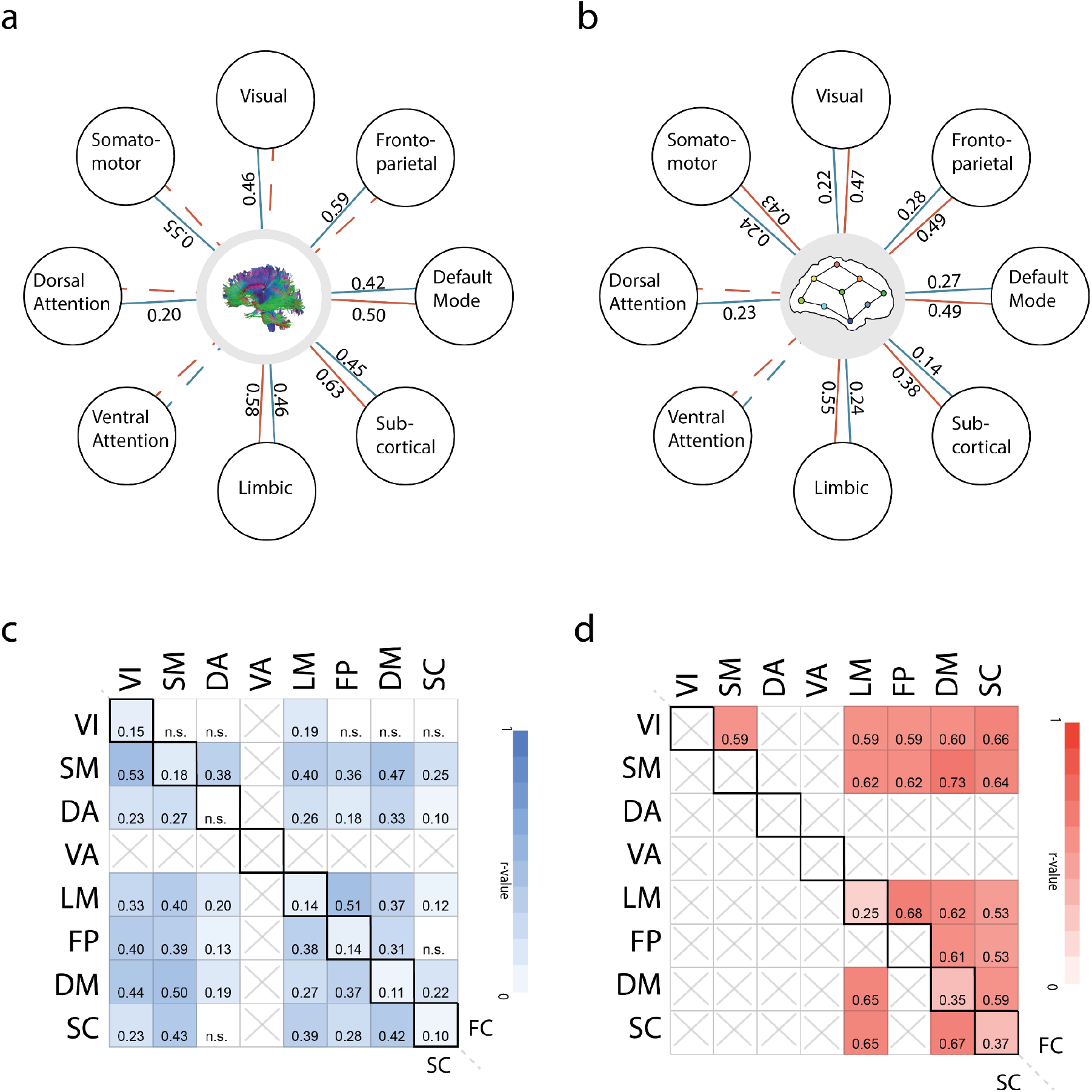
Brain network age prediction. (a) Structure and (b) Function age for canonical brain networks. Within-network connections for multiple networks successfully predicted postmenstrual age in term (blue lines) and preterm (red line) infants. Solid lines indicate significance at p<0.05, FDR-corrected, while dashed lines indicate non-significant predictions. **Correlations between predicted ages based on within-network connections for term (c) and preterm (d) infants**. Heatmaps show the correlation between predicted PMA from within-network connections. The upper triangle shows the correlation between functional ages. The lower triangle shows correlations between structural ages. The diagonal shows the correlations between structural and functional age for a network. (p<0.05; n.s. : not significant; box crossed: age not predictable from the within-network connections)

## Perinatal maternal exposures alter brain age gaps

Having established normative structural and functional ages, we asked how early exposures drive individual deviations from the normative developmental trajectories. In other words, how do prenatal and early postnatal exposures associate with structure and functional BAGs? BAGs were defined as predicted postmenstrual age minus real postmenstrual age. PMAs were regressed from the BAGs to adjust for the mean bias in the brain age prediction^29^. BAGs greater than 0 indicate that the brain looks older than it should be. BAGs less than 0 indicate the opposite. Maternal information was collected after birth. We studied ten prenatal and early postnatal measures across four common exposure types: maternal mental health (postnatal depression, lifetime psychiatry history), maternal physical health (age, BMI, high blood pressure, gestational diabetes), maternal demographics (age, education), and maternal substance use (smoking, alcohol). Linear models were used to associate BAGs and exposures.

For the term group, maternal age was positively correlated (r=0.13, p=0.0060) with structural BAGs, such that younger mothers had infants with younger structural connectomes. Postnatal depression was positively associated (t=2.11, p=0.036, df=374; r=0.11), such that mothers with higher EPDS scores had infants with older functional connectomes. Gestational diabetes was negatively associated with functional BAG (t=-2.11, p=0.038, df=435; r=-0.10), such that mothers with gestational diabetes had infants with younger functional connectomes. For preterm infants, maternal educational level was positively correlated with infant brain function age (r=0.23, p=0.012). Fig. 4 shows these associations. Overall, prenatal and early postnatal exposures influenced structural and functional brain ages.

**Fig. 4.**
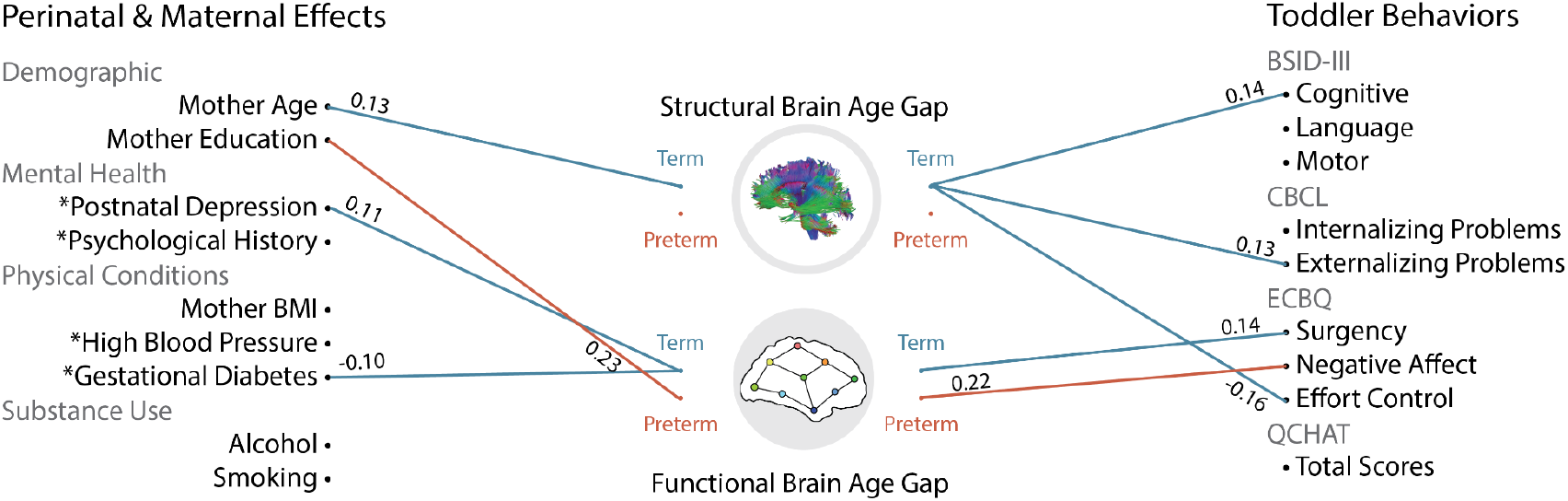
Brain age gaps (BAGs) associate with maternal effects and toddler behaviors. Maternal mental health, physical health, demographics, and substance use correlate with BAGs for term and preterm infants, controlling for the infant’s postmenstrual age. The BAGs also correlate with several later behaviors in toddlerhood, controlling for the infant’s postmenstrual age

## Brain age gaps associated with toddler behaviors

We further investigated if BAGs after birth exhibit a long-term influence on toddler behaviors. Cognitive and behavioral assessments were performed at 18 months. These assessments included the Bayley Scales of Infant and Toddler Development (BSID)^30^, the Child Behavior Checklist (CBCL)^31^, the Early Childhood Behavior Questionnaire (ECBQ)^32^, and the Quantitative Checklist for Autism in Toddlers (Q-CHAT)^33^. Table 1 shows the means and standard deviations for each. Correlations between the perinatal exposures and toddler outcomes are shown in Table S1-3. For term infants, structural BAGs were positively correlated with BSID cognitive (r=0.14, p=0.0077), CBCL externalizing problems (r=0.13, p=0.014), but negatively correlated with ECBQ effort control (r=-0.16, p=0.0025). These results indicate that the older structural connectomes at birth predict better cognitive function, worse emotion regulation, and worse externalizing problems. For term infants, functional BAGs were positively correlated with ECBQ surgency (r=0.14, p=0.0089). For preterm infants, the functional BAGs were positively correlated with ECBQ negative affect (r=0.22, p=0.01). These results indicate that the older functional connectomes at birth predict increased extroverted behaviors in term infants and worse mood in preterm infants. In exploratory analyses, we found that structural BAGs at birth partially mediated the association between maternal age and externalizing problems at 18 months old in the term group (average direct effect=-0.10, p<e-16; Total effect=-0.086, p=0.002; Prop. mediated=-0.14, p=0.030).

## Discussion

Using normative modeling, we establish that structural and functional connectomes accurately predict postmenstrual ages in term and preterm infants scanned within the first months of life. We observed greater predictive accuracy of structural connectomes compared to functional connectomes. Further, predicted structural and functional age were correlated. Preterm infants exhibited a greater correlation between structural and functional ages and weaker prediction from structural connectomes than term infants. At the network level, nearly all canonical brain networks—even putatively later developing ones—accurately predict PMA. In term infants, BAGs correlated with several early life exposures, including maternal depressive symptoms, age, and gestational diabetes. BAGs also correlated with toddler cognition, externalizing problems, extroverted behaviors, and emotion regulation. In preterm infants, BAGs correlated with maternal education and negative affect. These findings highlight the value of normative structural and functional development models in the perinatal period, where deviations from the normative trajectory are associated with early-life maternal exposures and later behavior.

Brain structural connectome develops before the functional connectome^4^. That structural connectomes consistently predicted PMA more accurately than functional connectomes for term infants aligns with this literature. Structural connectome is highly organized at birth with adult-like hubs and modules^2^. These richer features provide more information for the machine learning algorithms to learn from and increase prediction performance. Interestingly, between-network connections predicted PMA better than within-network connections, consistent with the rapid development of cross-hemisphere connections during this period^34^. The frontal lobe was the best predicting lobe for structural connectivity. Several core behaviors for infants are located in the frontal lobe. For example, morphometric features and structural connectivity in language regions (i.e. Broca’s region) are reportedly primed to learn language from birth^35,36^. The functional connectivity associated with these regions may take years to form^8,37–39^.

Nevertheless, PMA predictions with functional connectomes were highly accurate, with an average error of only 2.5 days worse than the structural connectomes. These results are consistent with recent research showing strong PMA prediction with functional connectomes in young infants^9^. In term infants, all networks predicted PMA (except ventral attention). Critically, there were no differences between higher-order networks that develop more slowly (e.g., DMN or FPN)^7^ and lower-order networks (e.g., motor or visual) that develop earlier^40,41^. Indeed, the FPN exhibited the greatest predictive accuracy. These results suggest that networks like the DMN or FPN might exhibit richer individual differences during the first postnatal month than previously thought. Additionally, the moderate correlation between structural and functional ages suggests a dependency between structural and functional development. Presumably, structural maturation serves as the foundation for later functional development. However, this causality cannot be inferred from our current results.

Preterm infants exhibited several different patterns in brain age compared to term infants. They had a higher correlation between structural and functional age and between structural and functional BAGs. In addition, structural connections at the individual network level were less predictive, while functional connections were more predictive in preterm than term infants (Fig 3, S1). Preterm birth accelerates functional trajectories due to earlier exposure to a postnatal environment^42^. This observation may explain the differences between term and preterm infants, such as the increased prediction accuracy from functional connectomes. Alternatively, preterm infants exhibit altered structural connectivity^43,44^—which, in turn, may be driving reduced prediction accuracy of the structural connectomes. A combination of altered structural and functional neurodevelopment is most likely responsible for these differences.

Several early life exposures affected brain age. Exposure to maternal distress (e.g., stress, depression, and anxiety) and substance use are commonly studied. Rather than focusing on brain regions of interest, such as the hippocampus and amygdala^45,46^, as in most works, our findings exhibit that altered connectivity extends beyond these regions and to the whole brain level. We also find altered structural and functional development for less commonly studied exposures like gestational diabetes and maternal age. Nevertheless, the biological pathways from exposures to altered brain development are unknown. Likely, multiple biological systems, including the maternal immune system and hypothalamic-pituitary-adrenal axis, are candidate pathways^47^. Many exposures activate inflammatory cytokines and proteins, which can then cross the placenta. It is hypothesized that while inflammatory cytokines contribute to fetal brain development, inappropriately high levels may alter cellular survival, proliferation, differentiation, axonal growth, and synaptogenesis^48,49^.

A hope for infant neuroimaging studies is to predict the risk for poor developmental outcomes in toddlerhood or later^50^. We found that BAGs are associated with several toddler outcomes. Importantly, these associations were not only in a single direction. In other words, younger and older connectomes were associated with worse outcomes. For example, a younger structural connectome is associated with worse cognition, while an older one is associated with worse externalizing problems and self-regulation. As the connectome putatively underlies cognition, a younger connectome may not support more advanced cognitive abilities. Conversely, early adverse caregiving can lead to accelerated brain development and increased risk of externalizing and regulation problems^51^. Delayed and accelerated development compared to one’s peers may be equally negative. This explanation aligns with the theory of critical periods in infancy. These periods are developmental windows where the brain is the most plastic to learning a skill or behavior. Changing the brain and the behavior it subserves outside this window is more difficult^52,53^. A normative range for BAGs could be used to define if an individual is missing a critical period. For example, a large BAG in either direction could suggest an infant is outside a critical period, resulting in worse developmental outcomes. However, longitudinal BAGs likely are needed to map when an individual enters and exits these critical periods and if they are indeed missed.

Our work has several strengths, including a large sample of infants with high-quality neuroimaging data, multimodal normative modeling, and longitudinal follow-up data in toddlerhood. However, several limitations should be acknowledged. First, while our study includes longitudinal data (prenatal exposures, postnatal MRI, and toddler behavior), measures were sparsely collected. Given the rapid change from infancy to toddlerhood, dense sampling is needed to account for the numerous interacting pre- and postnatal influences and establish direct relationships between prenatal exposures, brain development, and cognitive outcomes. It will also facilitate identifying critical periods of vulnerability and possible intervention. Second, the absence of genetic analysis in the current study might overlook the interplay between genetic factors and early life exposures on neurodevelopment. Genetic variations help shape individual differences in functional^54–56^ and structural connectivity^57–59^. These variations further interact with the environmental effects (i.e., epigenetics/maternal)^60–62^. Integrating genetic data into future studies could further nuance our results. Third, the reliance on maternal self-report measures and lack of detailed medical records may introduce bias and measurement errors. For example, we lack details about the frequency of tobacco and alcohol use and the actual blood pressure measurements. Replication of each early life exposure with detailed measurements during pregnancy is required.

In conclusion, we established normative models of structural and functional aging during the perinatal period. Results highlight the complex interplay between an infant’s brain age, early life exposures, and later cognitive behaviors. Overall, brain age and BAGs provide a valuable tool for studying neurodevelopment in infants.

## Methods

### Participants

All data were obtained from the Developing Human Connectome Project^28^ (dHCP, http://www.developingconnectome.org/), a longitudinal study of infant brain development with open access. The study received approval from the National Research Ethics Service West London committee. Participating families provided written consent before imaging. Our sample included 437 term infants (203 female, 234 male) and 174 preterm infants (74 female, 100 male) from the second data release of dHCP. Preterm infants were born between 23.00 and

36.86 weeks of gestation and scanned between 26.71 and 45.14 weeks. Term infants were born between 37.00 and 42.29 weeks of gestation and scanned between 37.43 and 44.71 weeks. Demographic information about the infants is presented in Table 1.

### MRI acquisition

All images were collected in the Evelina Newborn Imaging Centre, St Thomas’ Hospital, London, UK. MRI data were acquired with a Philips Achieva 3T scanner (Philips Medical Systems, Best, The Netherlands) with a dHCP-customized neonatal imaging system, including a 32-channel receive neonatal heal coil (Rapid Biomedical GmbH, Rimpar, DE)^28^. Infants were scanned during unsedated sleep after feeding and immobilization in a vacuum-evacuated bag, with hearing protection and physiological monitoring (including pulse oximetry, body temperature, and electrocardiography data) applied during scanning.

T2-weighted images were obtained using a Turbo spin echo sequence (TR=12s, TE=156ms, SENSE factor 2.11 (axial) and 2.54 (sagittal)) with overlapping slices (resolution = 0.8x0.8x1.6 mm3). T2w images were motion-corrected and super-resolved to a resolution of 0.8x0.8x0.8 mm3^63^. Diffusion-weighted imaging (DWI) was obtained in 300 directions (TR=3.8s, TE=90ms, SENSE factor 1.2, multiband factor 4, and resolution 1.5x1.5x3mm3 with 1.5 mm slice overlap) with b-values of 400s/mm2, 1000s/mm2 and 2600 s/mm2 spherically distributed in 64, 88 and 128 directions respectively using interleaved phase encoding^64^. BOLD fMRI runs were 15 minutes 3 seconds (2300 time volumes) and used a multislice gradient-echo planar imaging (EPI) sequence with TR 0.392s, TE 0.038s, flip angle 34 degrees, and acquired spatial resolution 2.15mm isotropic^65^.

### Structural connectome processing

Diffusion MRI was reconstructed at an in-plane resolution of 1.5mm and slice thickness of 1.5mm. Images were denoised and corrected for motion, eddy current, Gibbs ringing, and susceptibility artifact with the diffusion SHARD pipeline. In-scanner head motion was estimated by the SHARD outlier ratio, which is the mean outlier weight of all slices detected in slice-to-volume reconstruction. A quality check was conducted by neighboring DWI correction (NDC)^66^, leading to 34 scans being excluded due to their low NDC values calculated by median value-based outlier detection. The accuracy of b-table orientation was examined by comparing fiber orientations with those of a population-averaged template 56. Then, the reconstruction of the diffusion data was performed in native space with generalized q-sampling imaging (GQI)^67^ with a diffusion sampling length ratio of 1.25. The tensor metrics were calculated and analyzed using the resource allocation (TG-CIS200026) at Extreme Science and Engineering Discovery Environment (XSEDE) resources^68^.

After reconstructing images with GQI, the whole-brain fiber tracking was conducted with DSI-studio (http://dsi-studio.labsolver.org/) with quantitative anisotropy (QA) as the termination threshold. QA values were computed in each voxel in their native space for every subject. The tracking parameters were set as the angular cutoff of 60 degrees, step size of 1.0mm, minimum length of 30 mm, and maximum length of 300 mm. The whole-brain fiber tracking process was performed with the FACT algorithm until 1,000,000 streamlines were reconstructed for each individual. Here, we used a neonatal AAL-aligned brain parcellation with 90 nodes^69^ to construct the structural connectome for each infant. T2-weighted images in native DWI space were used to provide information on region segmentation during the construction of connectomes. The structural connectome for each individual was then constructed with a connectivity threshold of 0.001. The pairwise connectivity strength was calculated as the average QA value of each fiber connecting the two end regions. This resulted in a 90x90 adjacent matrix for each individual as the brain structural connectome matrix.

### Functional connectome processing

fMRI data was preprocessed with the dHCP functional pipeline^70^, including echo distortion correction, motion correction, independent component analysis (ICA) denoising, and registered to individual T2w native space. All subsequent preprocessing was performed with BioImage Suite (https://bioimagesuiteweb.github.io/webapp/). Mean time courses were regressed in white matter, cerebrospinal fluid, and gray matter. The neonatal AAL-aligned brain parcellation with 90 nodes was applied to the preprocessed fMRI data. Mean time courses of each node pair were correlated, and correlation coefficients were Fisher transformed, generating one 90x90 resting-state functional connectivity matrix per individual.

### Brain age

Predictive models of an infant’s PMA were built with connectome-based predictive modeling (CPM) using structural and functional connectomes. We used 10-fold cross-validation and a feature selection threshold of p=0.05^71^. Sex, brain volume, and head motion were controlled during feature selection. The predicted ages were then defined as the brain structure ages. The predictive models were trained in term and preterm infant groups separately. Brain age gap (BAG) was determined by subtracting the real postmenstrual age from the predicted postmenstrual age, serving as an indicator of whether the brain appears older (age gap>0) or younger (age gap<0) than expected. PMAs were regressed from the BAGs to adjust for the mean bias in the brain age prediction^29^.

To calculate network ages, connectomes were separated into eight canonical functional networks^72^: visual, somatomotor, frontoparietal, default mode, dorsal attention, ventral attention, limbic, and subcortical networks (see Table S4). For each network, the predicted PMA was generated based only on the connections within this network using the same CPM procedure described above. Similarly, we predict PMA using only the connections between two canonical networks. For validation purposes, we repeated the prediction process with support vector machines, and results were provided in supplementary information Table S5-7.

### Perinatal exposures

The influence of several perinatal exposures on infant brain development was investigated, as summarized in Table 2 and S8. Maternal physical health included body mass index (BMI), the occurrence of high blood pressure (i.e., pre-eclampsia, Hemolysis, Elevated Liver enzymes, and Low Platelets, or pregnancy-induced hypertension), and gestational diabetes. Maternal mental health included postnatal depression screened by the Edinburgh Postnatal Depression Scale (EPDS)^73^ and maternal history of mental health problems. Maternal substance use included the use of alcohol and tobacco during pregnancy. Maternal demographics included maternal age at expected delivery and education level. For the continuous measures (maternal age, BMI, EPDS level, alcohol, and smoking use level), Pearson’s correlations were used to quantify the association between the measures and infant brain age gaps. For the binary measures (psychological history, high blood pressure, gestational diabetes, and the use of recreational drugs), two-sample t-tests were used to test the effect on infant brains. The t-statistics were then transformed into r-values for comparison across measures.

**Table 2.** Maternal characteristics.

|  |  | n | mean (std) |
| --- | --- | --- | --- |
| Age at expected delivery |  | 611 | 33.80 (5.03) |
| Pre-pregnancy body mass index |  | 600 | 24.45 (4.60) |
| Last age in continuous education |  | 437 | 23.40 (4.99) |
| *Preeclampsia/eclampsia |  | 51/608 |  |
| *Gestational diabetes |  | 37/611 |  |
| *Psychological history |  | 79/600 |  |
| Postnatal depression |  | 535 | 5.51 (4.34) |
| Substance use during perinatal period | alcohol(number of units per week) | 45/606 | 2.56 (1.74) |
|  | smoking(number of cigarettes per day) | 19/606 | 9.05 (6.64) |

### Toddler outcomes

Four commonly-used behavioral measures were collected during the follow-up visits at 18 months (Table 1). Bayley Scales Of Infant and Toddler Development Third Version (BSID-III)^30^ evaluates three areas of development: cognitive, language, and motor. Child Behavior Checklist (CBCL)^31^ assesses the child’s psychological risks in internalizing problems, summing the anxious/depressed, withdrawn-depressed, and somatic complaints scores, and externalizing problems, including rule-breaking and aggressive behaviors. The Early Childhood Behavior Questionnaire (ECBQ) measures a toddler’s emotional and behavioral regulation skills through parents’ reports^32^. Quantitative Checklist for Autism in Toddlers (Q-CHAT) evaluates the risk in the social-emotional performance during toddlerhood^33^. Pearson’s correlations between these measures and BAGs were calculated.

## Data availability

Data from the Developing Human Connectome Project is publicly available (http://www.developingconnectome.org/).

## Code availability

The code is available at https://github.com/huiliii/infant_brain_age.

## Acknowledgement

This work is supported by NIH(R01MH126133). Data were provided by the developing Human Connectome Project, KCL-Imperial-Oxford Consortium funded by the European Research Council under the European Union Seventh Framework Programme (FP/2007-2013) / ERC Grant Agreement no. [319456]. We are grateful to the families who generously supported this trial.

## Author contributions

H.S.: conceptualization, methodology, investigation, result visualization, and writing-original draft. S.M., M.K. and X.H.: methodology, writing-review and editing; B.C. and M.S.: writing-review and editing; D.S.: conceptualization, supervision and writing-review and editing.

## Reference

1. Geng, X. et al. Structural and Maturational Covariance in Early Childhood Brain Development. Cerebral Cortex 27, 1795–1807 (2017).

2. Ball, G. et al. Rich-club organization of the newborn human brain. Proc. Natl. Acad. Sci. U.S.A. 111, 7456–7461 (2014).

3. Sun, H. et al. Network controllability of structural connectomes in the neonatal brain. Nat Commun 14, 5820 (2023).

4. Ciarrusta, J. et al. The developing brain structural and functional connectome fingerprint. Developmental Cognitive Neuroscience 55, 101117 (2022).

5. Gilmore, J. H., Knickmeyer, R. C. & Gao, W. Imaging structural and functional brain development in early childhood. Nat Rev Neurosci 19, 123–137 (2018).

6. Heuvel, M. I. van den & Thomason, M. E. Functional Connectivity of the Human Brain in Utero. Trends in Cognitive Sciences 20, 931–939 (2016).

7. Gao, W. et al. Functional Network Development During the First Year: Relative Sequence and Socioeconomic Correlations. Cerebral Cortex 25, 2919–2928 (2015).

8. Scheinost, D. et al. Functional connectivity for the language network in the developing brain: 30 weeks of gestation to 30 months of age. Cerebral Cortex 32, 3289–3301 (2022).

9. Nielsen, A. N. et al. Maturation of large-scale brain systems over the first month of life. Cerebral Cortex bhac242 (2022) doi:10.1093/cercor/bhac242.

10. Yin, C. et al. Anatomically interpretable deep learning of brain age captures domain-specific cognitive impairment. Proc. Natl. Acad. Sci. U.S.A. 120, e2214634120 (2023).

11. More, S. et al. Brain-age prediction: A systematic comparison of machine learning workflows. NeuroImage 270, 119947 (2023).

12. Kaufmann, T. et al. Common brain disorders are associated with heritable patterns of apparent aging of the brain. Nat Neurosci 22, 1617–1623 (2019).

13. Cole, J. H. et al. Brain age predicts mortality. Mol Psychiatry 23, 1385–1392 (2018).

14. Kardan, O. et al. Resting-state functional connectivity identifies individuals and predicts age in 8-to-26-month-olds. Developmental Cognitive Neuroscience 56, 101123 (2022).

15. Smyser, C. D. et al. Prediction of brain maturity in infants using machine-learning algorithms. NeuroImage 136, 1–9 (2016).

16. Li, Y. et al. Brain Connectivity Based Graph Convolutional Networks and Its Application to Infant Age Prediction. IEEE Transactions on Medical Imaging 41, 2764–2776 (2022).

17. Taoudi-Benchekroun, Y. et al. Predicting age and clinical risk from the neonatal connectome. NeuroImage 257, 119319 (2022).

18. Kawahara, J. et al. BrainNetCNN: Convolutional neural networks for brain networks; towards predicting neurodevelopment. NeuroImage 146, 1038–1049 (2017).

19. Kumpulainen, V. et al. Prenatal and Postnatal Maternal Depressive Symptoms Are Associated With White Matter Integrity in 5-Year-Olds in a Sex-Specific Manner. Biological Psychiatry 94, 924–935 (2023).

20. Qiu, A. et al. Maternal positive mental health during pregnancy impacts the hippocampus and functional brain networks in children. Nat. Mental Health 1–8 (2024) doi:10.1038/s44220-024-00202-8.

21. Lugo-Candelas, C. et al. Maternal Mental Health and Offspring Brain Development: An Umbrella Review of Prenatal Interventions. Biological Psychiatry 93, 934–941 (2023).

22. Rifkin-Graboi, A. et al. Antenatal Maternal Anxiety Predicts Variations in Neural Structures Implicated in Anxiety Disorders in Newborns. Journal of the American Academy of Child & Adolescent Psychiatry 54, 313-321.e2 (2015).

23. Gale-Grant, O. et al. Parental age effects on neonatal white matter development. NeuroImage: Clinical 27, 102283 (2020).

24. Kc, K., Shakya, S. & Zhang, H. Gestational Diabetes Mellitus and Macrosomia: A Literature Review. Annals of Nutrition and Metabolism 66, 14–20 (2015).

25. Magee, L. A. et al. Less-Tight versus Tight Control of Hypertension in Pregnancy. New England Journal of Medicine 372, 407–417 (2015).

26. Rompala, G., Nomura, Y. & Hurd, Y. L. Maternal cannabis use is associated with suppression of immune gene networks in placenta and increased anxiety phenotypes in offspring. Proceedings of the National Academy of Sciences 118, e2106115118 (2021).

27. Ross, E. J., Graham, D. L., Money, K. M. & Stanwood, G. D. Developmental Consequences of Fetal Exposure to Drugs: What We Know and What We Still Must Learn. Neuropsychopharmacol 40, 61–87 (2015).

28. Hughes, E. J. et al. A dedicated neonatal brain imaging system. Magnetic Resonance in Medicine 78, 794–804 (2017).

29. Smith, S. M., Vidaurre, D., Alfaro-Almagro, F., Nichols, T. E. & Miller, K. L. Estimation of brain age delta from brain imaging. NeuroImage 200, 528–539 (2019).

30. Bayley, N. Bayley Scales of Infant and Toddler Development, Third Edition. (2005) doi:10.1037/t14978-000.

31. Mazefsky, C. A., Anderson, R., Conner, C. M. & Minshew, N. Child Behavior Checklist Scores for School-Aged Children with Autism: Preliminary Evidence of Patterns Suggesting the Need for Referral. J Psychopathol Behav Assess 33, 31–37 (2011).

32. Putnam, S. P., Gartstein, M. A. & Rothbart, M. K. Measurement of fine-grained aspects of toddler temperament: The Early Childhood Behavior Questionnaire. Infant Behavior and Development 29, 386–401 (2006).

33. Allison, C. et al. The Q-CHAT (Quantitative CHecklist for Autism in Toddlers): A Normally Distributed Quantitative Measure of Autistic Traits at 18–24 Months of Age: Preliminary Report. J Autism Dev Disord 38, 1414–1425 (2008).

34. Thomason, M. E. et al. Cross-Hemispheric Functional Connectivity in the Human Fetal Brain. Sci. Transl. Med. 5, (2013).

35. Brauer, J., Anwander, A., Perani, D. & Friederici, A. D. Dorsal and ventral pathways in language development. Brain and Language 127, 289–295 (2013).

36. Witelson, S. F. & Pallie, W. LEFT HEMISPHERE SPECIALIZATION FOR LANGUAGE IN THE NEWBORN: NEUROANATOMICAL EVIDENCE OF ASYMMETRY. Brain 96, 641–646 (1973).

37. Emerson, R. W., Gao, W. & Lin, W. Longitudinal Study of the Emerging Functional Connectivity Asymmetry of Primary Language Regions during Infancy. J. Neurosci. 36, 10883–10892 (2016).

38. Perani, D. et al. Neural language networks at birth. Proceedings of the National Academy of Sciences 108, 16056–16061 (2011).

39. Bruchhage, M. M. K., Ngo, G.-C., Schneider, N., D’Sa, V. & Deoni, S. C. L. Functional connectivity correlates of infant and early childhood cognitive development. Brain Struct Funct 225, 669–681 (2020).

40. Allievi, A. G. et al. Maturation of Sensori-Motor Functional Responses in the Preterm Brain. Cerebral Cortex 26, 402–413 (2016).

41. Dehaene-Lambertz, G., Dehaene, S. & Hertz-Pannier, L. Functional Neuroimaging of Speech Perception in Infants. Science 298, 2013–2015 (2002).

42. Doria, V. et al. Emergence of resting state networks in the preterm human brain. Proceedings of the National Academy of Sciences 107, 20015–20020 (2010).

43. Ball, G. et al. Development of cortical microstructure in the preterm human brain. Proceedings of the National Academy of Sciences 110, 9541–9546 (2013).

44. Brenner, R. G., Wheelock, M. D., Neil, J. J. & Smyser, C. D. Structural and functional connectivity in premature neonates. Seminars in Perinatology 45, 151473 (2021).

45. Buss, C. et al. Maternal cortisol over the course of pregnancy and subsequent child amygdala and hippocampus volumes and affective problems. Proceedings of the National Academy of Sciences 109, E1312–E1319 (2012).

46. Qiu, A. et al. Maternal anxiety and infants’ hippocampal development: timing matters. Transl Psychiatry 3, e306–e306 (2013).

47. Monk, C., Lugo-Candelas, C. & Trumpff, C. Prenatal Developmental Origins of Future Psychopathology: Mechanisms and Pathways. Annual Review of Clinical Psychology 15, 317–344 (2019).

48. Mehler, M. F. & Kessler, J. A. Hematolymphopoietic and inflammatory cytokines in neural development. Trends in Neurosciences 20, 357–365 (1997).

49. Deverman, B. E. & Patterson, P. H. Cytokines and CNS Development. Neuron 64, 61–78 (2009).

50. Scheinost, D. et al. Machine Learning and Prediction in Fetal, Infant, and Toddler Neuroimaging: A Review and Primer. Biological Psychiatry 93, 893–904 (2023).

51. Luby, J. L. et al. Basic Environmental Supports for Positive Brain and Cognitive Development in the First Year of Life. JAMA Pediatrics (2024) doi:10.1001/jamapediatrics.2024.0143.

52. Margolis, E. T. & Gabard-Durnam, L. J. Prenatal influences on postnatal neuroplasticity: Integrating DOHaD and sensitive/critical period frameworks to understand biological embedding in early development. Infancy n/a,.

53. Nelson, C. A. & Gabard-Durnam, L. J. Early Adversity and Critical Periods: Neurodevelopmental Consequences of Violating the Expectable Environment. Trends in Neurosciences 43, 133–143 (2020).

54. Gao, W., Lin, W., Grewen, K. & Gilmore, J. H. Functional Connectivity of the Infant Human Brain: Plastic and Modifiable. Neuroscientist 23, 169–184 (2017).

55. Blanchett, R. et al. Genetic and environmental factors influencing neonatal resting-state functional connectivity. Cerebral Cortex 33, 4829–4843 (2023).

56. Stoecklein, S. et al. Variable functional connectivity architecture of the preterm human brain: Impact of developmental cortical expansion and maturation. Proceedings of the National Academy of Sciences 117, 1201–1206 (2020).

57. Knickmeyer, R. C. et al. Common Variants in Psychiatric Risk Genes Predict Brain Structure at Birth. Cerebral Cortex 24, 1230–1246 (2014).

58. Krishnan, M. L. et al. Machine learning shows association between genetic variability in PPARG and cerebral connectivity in preterm infants. Proceedings of the National Academy of Sciences 114, 13744–13749 (2017).

59. Alex, A. M. et al. Genetic Influences on the Developing Young Brain and Risk for Neuropsychiatric Disorders. Biological Psychiatry 93, 905–920 (2023).

60. Sparrow, S. et al. Epigenomic profiling of preterm infants reveals DNA methylation differences at sites associated with neural function. Transl Psychiatry 6, e716–e716 (2016).

61. Fumagalli, M. et al. From early stress to 12-month development in very preterm infants: Preliminary findings on epigenetic mechanisms and brain growth. PLOS ONE 13, e0190602 (2018).

62. Qiu, A. et al. COMT Haplotypes Modulate Associations of Antenatal Maternal Anxiety and Neonatal Cortical Morphology. AJP 172, 163–172 (2015).

63. Cordero-Grande, L., Hughes, E. J., Hutter, J., Price, A. N. & Hajnal, J. V. Three-dimensional motion corrected sensitivity encoding reconstruction for multi-shot multi-slice MRI: Application to neonatal brain imaging. Magnetic Resonance in Medicine 79, 1365–1376 (2018).

64. Scheid, B. H. et al. Time-evolving controllability of effective connectivity networks during seizure progression. Proceedings of the National Academy of Sciences 118, e2006436118 (2021).

65. Price, A. N. et al. Accelerated Neonatal fMRI using Multiband EPI.

66. Yeh, F.-C. et al. Differential tractography as a track-based biomarker for neuronal injury. NeuroImage 202, 116131 (2019).

67. Yeh, F.-C., Wedeen, V. J. & Tseng, W.-Y. I. Generalized $ q$-Sampling Imaging. IEEE Transactions on Medical Imaging 29, 1626–1635 (2010).

68. Towns, J. et al. XSEDE: Accelerating Scientific Discovery. Comput. Sci. Eng. 16, 62–74 (2014).

69. Shi, F. et al. Infant Brain Atlases from Neonates to 1- and 2-Year-Olds. PLOS ONE 6, e18746 (2011).

70. Fitzgibbon, S. P. et al. The developing Human Connectome Project (dHCP) automated resting-state functional processing framework for newborn infants. NeuroImage 223, 117303 (2020).

71. Shen, X. et al. Using connectome-based predictive modeling to predict individual behavior from brain connectivity. Nat Protoc 12, 506–518 (2017).

72. Thomas Yeo, B. T. et al. The organization of the human cerebral cortex estimated by intrinsic functional connectivity. J Neurophysiol 106, 1125–1165 (2011).

73. Matthey, S., Barnett, B. & White, T. The Edinburgh Postnatal Depression Scale. The British Journal of Psychiatry 182, 368–368 (2003).

