## Supplementary Information for "Brain age prediction and deviations from normative trajectories in the neonatal connectome"

**Associations between maternal conditions and infant behaviors during toddlerhood**

Table S1. Correlations (r-value) between maternal conditions and toddler behaviors in the whole group

| Maternal Conditions | BSID | | | CBCL | | | ECBQ | | | QCHAT |
|---|---|---|---|---|---|---|---|---|---|---|
| | Cognitive | Language | Motor | Internalizing | Externalizing | Total | Surgency | Negative Affect | Effort Control | Total Score |
| Postnatal Depression | -0.10* | -0.14** | | 0.23*** | 0.24*** | 0.27*** | | 0.17*** | -0.14** | 0.16*** |
| *Psychological History | | | | | 0.11* | | | | | |
| Age | | | | -0.16*** | -0.10* | -0.14** | | | | -0.11* |
| BMI | -0.19*** | -0.13** | -0.16*** | | | | 0.09* | 0.16*** | | 0.13** |
| *High Blood Pressure | | | | | | | | | | |
| *Gestational Diabetes | | | | | | | | | | |
| Education | 0.10* | 0.11* | | | | | | | | |
| Alcohol | | 0.11* | 0.12* | | | -0.11* | | | | |
| Smoking | | -0.10* | 0 | 0.16*** | 0.14** | 0.17*** | | 0.11* | | 0.14** |

**:p<0.05,**:p<0.01,***:p<0.001*

Table S2. Correlations (r-value) between maternal conditions and toddler behaviors in the term group

| Maternal Conditions | BSID | | | CBCL | | | ECBQ | | | QCHAT |
|---|---|---|---|---|---|---|---|---|---|---|
| | Cognitive | Language | Motor | Internalizing | Externalizing | Total | Surgency | Negative Affect | Effort Control | Total Score |
| Postnatal Depression | | -0.12* | | 0.19** | 0.27*** | 0.27*** | 0.16** | -0.12* | | |
| *Psychological History | | | | | | | | | | |
| Age | | 0.13* | | -0.23*** | | -0.17** | | | -0.16** | |
| BMI | -0.15** | | -0.12* | | | | 0.11 | | | |
| *High Blood Pressure | | | | | | | | | | |
| *Gestational Diabetes | | | | | | | | | | |
| Education | | 0.12* | | | | | | | | |
| Alcohol | | 0.18*** | 0.11* | | | -0.11* | | | | |
| Smoking | | | | | | | | | | |

**:p<0.05,**:p<0.01,***:p<0.001*

Table S3. Correlations (r-value) between maternal conditions and toddler behaviors in the preterm group

| Maternal conditions | BSID | | | CBCL | | | ECBQ | | | QCHAT |
|---|---|---|---|---|---|---|---|---|---|---|
| | Cognitive | Language | Motor | Internalizing | Externalizing | Total | Surgency | Negative Affect | Effort Control | Total Score |
| Postnatal Depression | -0.20* | | | 0.29** | | 0.26** | | 0.21** | | 0.26** |
| *Psychological History | | | | | 0.19* | | | | | |
| Age | | | | | | | | | | |
| BMI | -0.27* | -0.21* | -0.24* | | | | | 0.27** | | 0.22* |
| *High Blood Pressure | | | | | | | | | | |
| *Gestational Diabetes | | | | | | | | | | |
| Education | | | | | | | | | | |
| Alcohol | | | | | | | | | | |
| Smoking | | | | 0.26** | 0.24** | 0.29*** | | 0.22* | | 0.24** |

**:p<0.05,**:p<0.01,***:p<0.001*

Table S4. Neonatal 90 node parcellation's regional names and resting-state networks(RSNs)

| Node | Region | RSNs | Node | Region | RSNs |
|---|---|---|---|---|---|
| 1 | Precentral gyrus left | Dorsal Attention | 46 | Cuneus right | Subcortical |
| 2 | Precentral gyrus right | Dorsal Attention | 47 | Lingual gyrus left | Visual |
| 3 | Superior frontal gyrus (dorsal) left | Dorsal Attention | 48 | Lingual gyrus right | Visual |
| 4 | Superior frontal gyrus (dorsal) right | Dorsal Attention | 49 | Superior occipital gyrus left | Visual |
| 5 | Orbitofrontal cortex (superior) left | Frontoparietal | 50 | Superior occipital gyrus right | Visual |
| 6 | Orbitofrontal cortex (superior) right | Frontoparietal | 51 | Middle occipital gyrus left | Visual |
| 7 | Middle frontal gyrus left | Frontoparietal | 52 | Middle occipital gyrus right | Visual |
| 8 | Middle frontal gyrus right | Frontoparietal | 53 | Inferior occipital gyrus left | Visual |
| 9 | Orbitofrontal cortex (middle) left | Frontoparietal | 54 | Inferior occipital gyrus right | Visual |
| 10 | Orbitofrontal cortex (middle) right | Frontoparietal | 55 | Fusiform gyrus left | Visual |
| 11 | Inferior frontal gyrus (opercular) left | Default Mode | 56 | Fusiform gyrus right | Visual |
| 12 | Inferior frontal gyrus (opercular) right | Default Mode | 57 | Postcentral gyrus left | Visual |
| 13 | Inferior frontal gyrus (triangular) left | Somatomotor | 58 | Postcentral gyrus right | Visual |
| 14 | Inferior frontal gyrus (triangular) right | Somatomotor | 59 | Superior parietal gyrus left | Visual |
| 15 | Orbitofrontal cortex (inferior) left | Somatomotor | 60 | Superior parietal gyrus right | Visual |
| 16 | Orbitofrontal cortex (inferior) right | Somatomotor | 61 | Inferior parietal lobule left | Somatomotor |
| 17 | Rolandic operculum left | Limbic | 62 | Inferior parietal lobule right | Somatomotor |
| 18 | Rolandic operculum right | Limbic | 63 | Supramarginal gyrus left | Dorsal Attention |
| 19 | Supplementary motor area left | Default Mode | 64 | Supramarginal gyrus right | Dorsal Attention |
| 20 | Supplementary motor area right | Default Mode | 65 | Angular gyrus left | Frontoparietal |
| 21 | Olfactory left | Default Mode | 66 | Angular gyrus right | Frontoparietal |
| 22 | Olfactory right | Default Mode | 67 | Precuneus left | Somatomotor |
| 23 | Superior frontal gyrus (medial) left | Limbic | 68 | Precuneus right | Somatomotor |
| 24 | Superior frontal gyrus (medial) right | Limbic | 69 | Paracentral lobule left | Frontoparietal |

| | | | | | |
|---|---|---|---|---|---|
| 25 | Orbitofrontal cortex (medial) left | Limbic | 70 | Paracentral lobule right | Frontoparietal |
| 26 | Orbitofrontal cortex (medial) right | Limbic | 71 | Caudate left | Default Mode |
| 27 | Rectus gyrus left | Limbic | 72 | Caudate right | Default Mode |
| 28 | Rectus gyrus right | Limbic | 73 | Putamen left | Somatomotor |
| 29 | Insula left | Limbic | 74 | Putamen right | Somatomotor |
| 30 | Insula right | Limbic | 75 | Pallidum left | Subcortical |
| 31 | Anterior cingulate gyrus left | Limbic | 76 | Pallidum right | Subcortical |
| 32 | Anterior cingulate gyrus right | Limbic | 77 | Thalamus left | Subcortical |
| 33 | Middle cingulate gyrus left | Ventral Attention | 78 | Thalamus right | Subcortical |
| 34 | Middle cingulate gyrus right | Ventral Attention | 79 | Heschl gyrus left | Subcortical |
| 35 | Posterior cingulate gyrus left | Default Mode | 80 | Heschl gyrus right | Subcortical |
| 36 | Posterior cingulate gyrus right | Default Mode | 81 | Superior temporal gyrus left | Subcortical |
| 37 | Hippocampus left | Ventral Attention | 82 | Superior temporal gyrus right | Subcortical |
| 38 | Hippocampus right | Ventral Attention | 83 | Temporal pole (superior) left | Somatomotor |
| 39 | ParaHippocampal gyrus left | Default Mode | 84 | Temporal pole (superior) right | Somatomotor |
| 40 | ParaHippocampal gyrus right | Default Mode | 85 | Middle temporal gyrus left | Somatomotor |
| 41 | Amygdala left | Subcortical | 86 | Middle temporal gyrus right | Somatomotor |
| 42 | Amygdala right | Subcortical | 87 | Temporal pole (middle) left | Limbic |
| 43 | Calcarine cortex left | Subcortical | 88 | Temporal pole (middle) right | Limbic |
| 44 | Calcarine cortex right | Subcortical | 89 | Inferior temporal gyrus left | Default Mode |
| 45 | Cuneus left | Subcortical | 90 | Inferior temporal gyrus right | Default Mode |

### Age prediction with inter-network connections

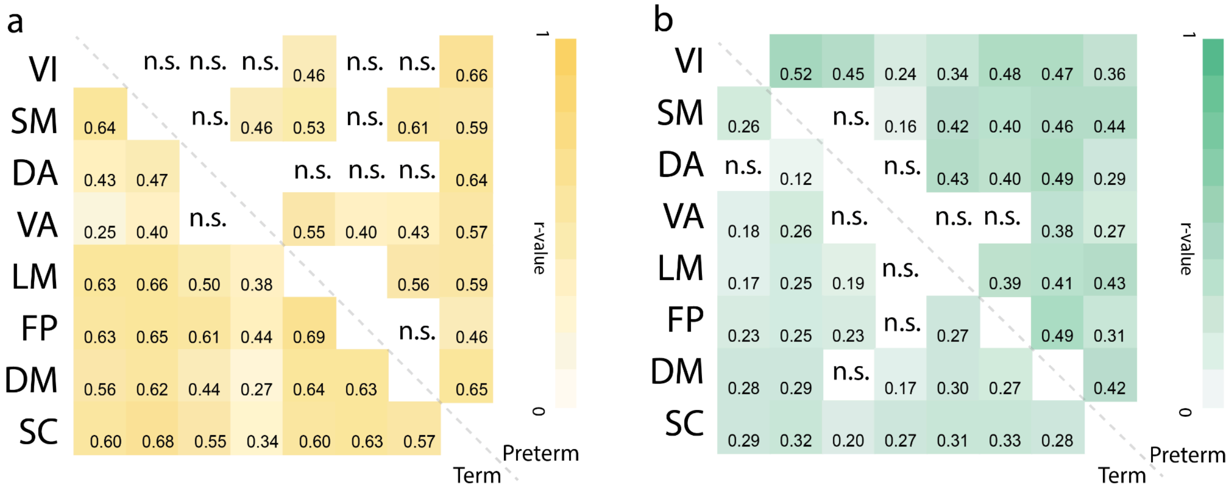


**Fig. S1. PMA prediction with the between network connections.** Multiple between-network **(a)** structural and **(b)** functional connections accurately predicted postmenstrual ages for term (lower triangular of the heat maps) and preterm (upper triangular of the heat maps) infants.

**Brain lobe age**

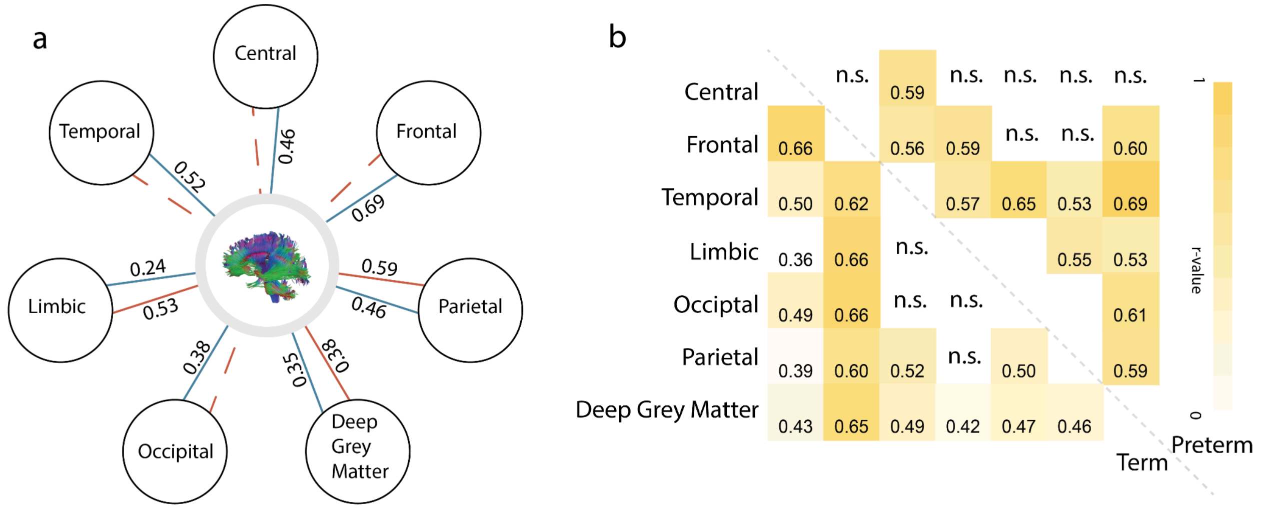


**Fig.S2. Brain lobe age prediction. (a) Structure age for each brain lobe.** Divided anatomically into seven areas: central, frontal, parietal, temporal, limbic, occipital lobes, and deep gray matter. PMA could be predicted for term infants (as blue lines) based on the white-matter connections within each lobe. For preterm infants (as red lines), only connections in the parietal and limbic lobes and deep gray matter could predict their PMA. **(b) Age prediction with inter-lobe structural connections.** Multiple inter-lobe structural connections managed to predict the real infant ages, with the heatmap showing the correlations between the predicted and real ages (upper triangular for preterm infants and lower triangular for term infants).

### Replication with SVM to predict brain ages

Table S5. Age prediction with SVM

| | term | | | | preterm | | | |
|---|---|---|---|---|---|---|---|---|
| | structure | | function | | structure | | function | |
| | *r* | MAE(week) | *r* | MAE(week) | *r* | MAE(week) | *r* | MAE(week) |
| whole-brain | 0.77*** | 0.88 | 0.51*** | 1.23 | 0.76*** | 1.54 | 0.54*** | 2.3 |
| visual | 0.54*** | 1.18 | 0.32*** | 1.39 | 0.65 | 2.00 | 0.49*** | 2.2958 |
| somatomotor | 0.55*** | 1.18 | 0.26*** | 1.42 | 0.53 | 2.26 | 0.51*** | 2.3898 |
| dorsal attention | 0.36*** | 1.34 | 0.24*** | 1.37 | 0.70 | 2.19 | 0.46*** | 2.3289 |
| ventral attention | n.s. | | n.s. | | n.s. | | 0.20** | 2.5596 |
| limbic | 0.50*** | 1.24 | 0.33*** | 1.36 | 0.56 | 2.12 | 0.52*** | 2.3591 |
| frontoparietal | 0.61*** | 1.12 | 0.24*** | 1.40 | 0.63 | 2.19 | 0.54*** | 2.3203 |
| default mode | 0.46*** | 1.26 | 0.34*** | 1.35 | 0.56 | 2.22 | 0.44*** | 2.4105 |
| subcortical | 0.50*** | 1.23 | 0.38*** | 1.30 | 0.61 | 2.27 | 0.45*** | 2.3861 |

**:p<0.05,**:p<0.01,***:p<0.001; n.s: not significant*

Table S6. Association between BAGs predicted with SVM and early life exposures

| | term | | | | preterm | | | |
|---|---|---|---|---|---|---|---|---|
| | structure | | function | | structure | | function | |
| | *r* | *p* | *r* | *p* | *r* | *p* | *r* | *p* |
| Postnatal Depression | 0.00 | 0.98 | -0.02 | 0.71 | -0.10 | 0.22 | 0.00 | 0.99 |
| *Psychological History | -0.05 | 0.26 | 0.06 | 0.25 | -0.15 | 0.05 | -0.12 | 0.13 |
| Age | **-0.11** | **0.03** | -0.04 | 0.42 | 0.03 | 0.69 | 0.04 | 0.58 |
| BMI | 0.02 | 0.69 | -0.03 | 0.56 | -0.05 | 0.55 | 0.05 | 0.54 |
| *High Blood Pressure | 0.00 | 0.93 | 0.05 | 0.34 | -0.11 | 0.14 | -0.16 | 0.04 |
| *Gestational Diabetes | -0.03 | 0.48 | **0.03** | **0.52** | -0.07 | 0.37 | -0.01 | 0.86 |
| Education | -0.08 | 0.18 | 0.01 | 0.90 | 0.11 | 0.23 | **-0.09** | **0.35** |
| Alcohol | -0.02 | 0.63 | 0.01 | 0.91 | 0.01 | 0.88 | -0.18 | 0.02 |
| Smoking | 0.05 | 0.33 | -0.18 | 0.00 | -0.03 | 0.72 | -0.02 | 0.80 |

Table S7. Association between BAGs predicted with SVM and toddler behaviors

| | term | | | | preterm | | | |
|---|---|---|---|---|---|---|---|---|
| | structure | | function | | structure | | function | |
| | *r* | *p* | *r* | *p* | *r* | *p* | *r* | *p* |
| *BSID* | | | | | | | | |
| Cognition | **-0.13** | **0.01** | 0.00 | 0.94 | 0.00 | 0.99 | 0.02 | 0.80 |
| Language | **-0.11** | **0.05** | 0.04 | 0.46 | -0.10 | 0.28 | 0.10 | 0.26 |
| Motor | 0.02 | 0.67 | -0.03 | 0.53 | -0.04 | 0.67 | -0.03 | 0.73 |
| *CBCL* | | | | | | | | |
| Internalizing | 0.05 | 0.34 | 0.06 | 0.27 | 0.11 | 0.21 | -0.08 | 0.37 |
| Externalizing | **0.09** | **0.10** | 0.07 | 0.19 | 0.05 | 0.61 | 0.09 | 0.31 |
| Total | **0.10** | **0.08** | 0.07 | 0.21 | 0.08 | 0.37 | 0.02 | 0.79 |
| *ECBQ* | | | | | | | | |
| Surgency | 0.10 | 0.08 | **0.10** | **0.07** | -0.03 | 0.76 | 0.15 | 0.11 |
| Negative Affect | 0.06 | 0.26 | 0.01 | 0.84 | 0.02 | 0.83 | **-0.20** | **0.02** |
| Effort Control | **0.00** | **0.96** | -0.04 | 0.41 | -0.05 | 0.61 | -0.07 | 0.44 |
| *Q-CHAT* | | | | | | | | |
| Total | 0.09 | 0.09 | 0.04 | 0.48 | 0.10 | 0.25 | -0.12 | 0.18 |

Table S8. Maternal characteristics by birth types

| | | Preterm | | Term | | |
|---|---|---|---|---|---|---|
| | | n | mean (std) | n | mean (std) | *p* Value |
| Age at expected delivery | | 174 | 34.22 (5.56) | 437 | 33.63 (4.81) | .19 |
| Pre-pregnancy body mass index | | 170 | 24.70 (4.83) | 430 | 24.34 (4.51) | .39 |
| Last age in continuous education | | 119 | 22.76 (4.62) | 318 | 23.64 (5.10) | .10 |
| *Preeclampsia/eclampsia | | 29/174 | | 22/434 | | <0.001 |
| *Gestational diabetes | | 14/174 | | 23/437 | | .19 |
| *Psychological history | | 21/170 | | 58/430 | | .71 |
| Postnatal depression | | 159 | 6.45 (4.93) | 376 | 5.11 (4.01) | <0.01 |
| Substance use during perinatal period | alcohol | 13/170 | 2.54 (1.27) | 436 | 2.56 (1.92) | .94 |
| | smoking | 10/170 | 10.60 (8.41) | 9/436 | 7.33 (3.64) | <0.01 |
